# RetroFun-RVS: a retrospective family-based framework for rare variant analysis incorporating functional annotations

**DOI:** 10.1101/2022.06.21.497085

**Authors:** Loïc Mangnier, Ingo Ruczinski, Jasmin Ricard, Claudia Moreau, Simon Girard, Michel Maziade, Alexandre Bureau

## Abstract

A large proportion of genetic variations involved in complex diseases are rare and located within non-coding regions, making the interpretation of underlying biological mechanisms a daunting task. Although technical and methodological progresses have been made to annotate the genome, current disease - rare-variant association tests incorporating such annotations suffer from two major limitations. Firstly, they are generally restricted to case-control designs of unrelated individuals, which often require tens or hundreds of thousands of individuals to achieve sufficient power. Secondly, they were not evaluated with region-based annotations needed to interpret the causal regulatory mechanisms. In this work we propose RetroFun-RVS, a new retrospective family-based score test, incorporating functional annotations. One of the critical features of the proposed method is to aggregate genotypes while measuring rare variant sharing among affected family members to compute the test statistic. Through extensive simulations, we have demonstrated that RetroFun-RVS integrating networks based on 3D genome contacts as functional annotations reaches greater power over the region-wide test, other strategies to include sub-regions and competing methods. Also, the proposed framework shows robustness to non-informative annotations, keeping a stable power when causal variants are spread across regions. We provide recommendations when dealing with different types of annotations or family structures commonly encountered in practice. Application of RetroFun-RVS is illustrated on whole genome sequence in the Eastern Quebec Schizophrenia and Bipolar Disorder Kindred Study with networks constructed from 3D contacts and epigenetic data on neurons. In summary we argue that RetroFun-RVS, by allowing integration of functional annotations corresponding to regions or networks with transcriptional impacts, is a useful framework to highlight regulatory mechanisms involved in complex diseases.

## 1 Introduction

Over the past few years with the democratization of whole-exome or whole-genome sequencing data, important progresses have been made in the effort to link genetic variations to phenotypes. Indeed, at population scale, Genome-Wide Association Studies (GWAS) have provided useful resources to highlight variants involved in diseases. However, these methods, in addition to requiring tens or hundreds of thousands of individuals, are mainly restricted to common variants, leaving an important part of heritability unexplained ^1^. In fact, studies have shown that the individual genetic risk is also substantially influenced by rare variants (minor allele frequency (MAF) *≤* 1%), ^2,3^. In addition to being rare, variants influencing disease risk tend to be located within non-coding regions, making the underlying biological mechanisms difficult to interpret ^4^. Thus, the tremendous amount of rare variants located within non-coding regions brings new challenges to identify new causal variants involved in diseases, and accounting for their functional impacts remains crucial from a fine-mapping perspective, hence translational medicine applications ^5^.

Methods have been proposed to overcome the challenge of sparsity. Indeed, because variants are rare, methods testing them in an unitary fashion perform badly ^6^. Thus, rare-variants association tests (RVATs) are methods aggregating genotypes across several variant sites within a gene, pathway or regions functionally close. By collapsing variants over regions, these methods considerably reduce the number of tests throughout the genome, hence increasing statistical power. Among them, burden tests were initially proposed and are powerful when all variants across regions show a homogeneous effect ^7,6^. However, when regions combine both deleterious and protective variants, burden tests comparing cases to controls suffer from a substantial decrease of power. Alternatives to address this limitation have been proposed ^8,9^. One of the critical features of RVATs is that they can be expressed through regression models, allowing either the integration of covariates or variant weights, either fixed (based on the MAF), or estimated in a data adaptive manner ^6,10^.

An alternative approach is to exploit family-based studies. In addition to reducing genetic heterogeneity, pedigree-based studies have been shown to have more power than population-based approaches for detecting rare variants, when an enrichment of risk variants among families is expected ^11,12,13^. Information provided by variants segregating with the disease, even imperfect, can be exploited to highlight new causal variants, giving a second breath to studies in extended pedigrees ^14^. Recent methods based on identity-by-descent (IBD) or combining both linkage approaches and RVATs have been developed ^15,16,17^. These approaches focus on, or can be restricted to only affected family members, when these are expected to contribute more information than unaffected subjects^18^. Affected-only designs have a long tradition in gene-gene or gene-environment interaction analysis and have been extended to family-based studies, requiring smaller sample sizes to reach equivalent power, compared to considering unrelated case-only individuals, which is an appealing feature in practice ^19^. However, in many cases, knowing and defining the sampling scheme is difficult, hence impossible, pushing researchers to consider retrospective approaches. Retrospective models by conditioning on phenotypes do not explicitly model the ascertainment process. Successful applications of such methods have been shown for common ^18^ and rare variants ^20^.

A limitation of all the above methods is that none of them currently integrates external information on biological mechanisms involved in diseases. How to leverage information on non-coding regulatory elements in the detection of variants influencing disease risk remains an open question. Thus, there is an increasing interest in using external information for this task, and hence highlighting the biological mechanisms. Recent methods, such as FST ^21^ or FunSPU ^22^ have proposed to adaptively test functional annotations under a general RVAT framework. These methods have shown substantial increases in power when at least one functional score is predictive for the effect of variants on the trait, while they show robustness when no annotations were predictive for variant impact on the trait, revealing new causal variants involved in complex traits. Moreover, the multiple ways to define test statistics corresponding to several functional annotations created a need for combining p-values within a given region to assess the association with a trait, while adjusting for multiplicity. Liu et al. ^23^ have proposed the aggregated Cauchy association test (ACAT), a powerful statistical framework combining p-values in an efficient way, not requiring resampling procedures, nor independent p-values nor explicit models for correlations. This facilitating applications even at the genome-wide scale. Although these set-based tests have made possible the discovery of new regions involved in complex diseases, they required very large sample sizes of unrelated subjects.

More recently, with the striking development of methods detecting regulatory elements such as enhancers ^24,25^, progresses have been made in associating non-coding SNPs to their target genes ^26,27^. Subsequently, some authors have proposed to incorporate this information within statistical frameworks. Ma et al. ^28^ have demonstrated that long range 3D interactions between genes and enhancers add information for the integration of non-coding regulatory regions within gene-based frameworks. This model, consistent with previous studies, only considers pairs of gene-enhancer, ^29^. Frameworks extending gene-enhancer pairs to Cis-Regulatory Hubs (CRHs), networks encompassing up to several genes and active enhancers have been proposed ^30^. CRHs have been shown to be a relevant model in schizophrenia etiology, explaining more heritability than tissue- and non-tissue-specific elements, and being more effective to link noncoding SNPs to differentially expressed genes in schizophrenia compared to Topologically Associated Domains (TADs) or pairs of gene-enhancer. To our knowledge, no study to date has proposed to integrate functional annotations within a family-based RVAT framework, while allowing the incorporation of discontinuous genomic regions involved in 3D-based networks.

In this paper, we propose RetroFun-RVS (Retrospective Functional Rare Variant Sharing), a model, allowing the integration of functional annotations under a family-based design considering only affected individuals. Through extensive simulation studies, we have demonstrated that RetroFun-RVS integrating CRHs as functional annotations is a more powerful approach to detect causal variants over other strategies, while well controlling the Type I error rate. We provide recommendations when dealing with different types of functional scores or pedigree structures. Finally, illustrating RetroFun-RVS on the whole genome sequence in the Eastern Quebec Schizophrenia and Bipolar disorder Kindred study we have demonstrated that integrating 3D-based functional annotations through networks is a relevant strategy to gain power for detection of causal variants, while highlighting the underlying biological mechanisms involved in diseases.

## 2 Material and Methods

### 2.1 Notations and Model

Suppose that we have *N* subjects within *F* families, where *n*_*f*_ is the number of individuals for the *f*^*th*^ family. Let’s define *Y*, a binary vector of phenotypes, *G* a *N × p* matrix of genotypes for rare variants, coded as unordered, discrete variables. Assuming a log-additive model for the individual SNP effect on disease risk, under the assumptions of rare disease for all genotypes (i.e. weak variant penetrance) and of conditional independence of the phenotypes of different individuals given their genotypes and considering only affected individuals, following Schaid et al. ^18^, the retrospective likelihood for one family can be written as:

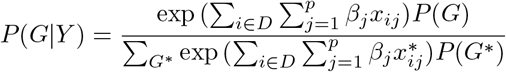

where *D* is the subset of affected members in the family, while *x*_*ij*_ is a condensed notation for *x*(*G*_*ij*_), the number of minor alleles {0, 1, 2} for variant *j* in individual *i* in the multilocus genotype configuration *G* for all family members. Also, we assume that only one copy of the minor allele was introduced once by a family founder, implying *x*_*ij*_ can only take the values 0 or 1 in the absence of inbreeding in the family (occasional genotypes with 2 variant alleles may be recoded as *x*_*ij*_ = 1 with little impact on the results). In presence of inbreeding and/or cryptic relatedness among family founders, homozygous genotypes for rare alleles are expected and are allowed in the RetroFun-RVS implementation using options described in the Supplementary Material and Methods.

In Schaid et al. ^18^, *P* (*G*) is the unconditional genotype probability and depends on MAF, which needs to be estimated in practice. However, obtaining accurate estimates of rare variant MAFs in a population is difficult. Instead, we opted for conditioning the probability on the event of observing at least one a copy of each RV *j* present in the family (i.e., ∑_*i*_ *x*_*ij*_ *≥* 1) as in ^15^. In addition, we combined this conditional probability with the assumption that the variant frequency tends to 0, hence the probability does not depend on MAF and therefore the computation does not require external variant frequency estimates. In this context, the genotypes can be interpreted as rare variant sharing patterns (referring to as RVS in the method name). The sum in the denominator is over all genotype configurations respecting the condition within the given pedigree, where *G*^*∗*^ denotes one particular configuration. Since we expect that risk variant effects dominate protective variant effects in the score test statistic when considering only affected individuals (Supplementary Materials and Methods and Figure S1), we propose to adapt the retrospective framework for a burden test ^7,6^. Hence, we can express *β*_*j*_ the effect of the *j*^*th*^ variant through *w*_*j*_*β*_0_ where *w*_*j*_ is usually a weighting function to specify variant effects through a function of MAF.

As suggested by He et al. ^21^, the effect for the *j*^*th*^ variant can be partitioned into effect parameters *γ*_*k*_ with respect to functional annotations *Z*_*jk*_, *k* = 1 … *q*. Consequently, under a burden test framework this leads to:

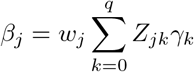

with *Z*_*j*0_ = 1 and *γ*_0_ corresponding to the original burden test parameter. Intuitively, this partition of the variant effect allows a modulation of the variant effect based on MAF and functional annotations. Moreover, when no predictive functional annotations are present for the trait, the burden of all *p* variants may nonetheless capture an overall effect on risk, and testing *γ*_0_ ensures the combined test has some power. When at least one annotation is predictive, the partitioned model offers increased power over the original test ^21^.

Now combining the retrospective likelihood model described by ^18^ and the decomposed variant effect, we obtain:

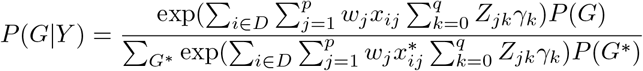

Thus for the *k*^*th*^ functional annotation the score function *S*_*k*_ summed across the *F* families is :

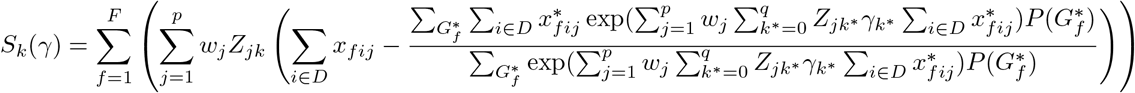

Intuitively, this quantity can be seen as the difference between the observed genotype value and the expected value, weighted by MAF and functional annotations. Setting *γ* to 0, we obtain the score statistic for the *k*^*th*^ functional annotation:

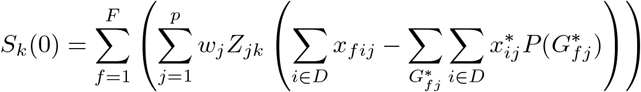

The genotype probability required *P* (*G*_*fj*_) is for a single variant configuration in family *f* and can be computed using RVS ^31^. *Q*_*k*_ is the test statistic corresponding to *S*_*k*_(0), asymptotically following a normal distribution with mean 0 and variance obtained by combining sharing pattern probabilities under the null and observed genotypes within families. Moreover, simplifications may be obtained from assumptions on the linkage disequilibrium structure (See Supplementary Materials and Methods). However, we observed when only few variants are expected within a functional annotation or a small number of families is observed that resampling procedures may be required to adequately control the Type I error rate (See next sub-section Bootstrap procedure using rare variant sharing patterns).

For testing multiple functional scores within a single unified test *H*_0_ : *∀k, γ*_*k*_ = 0 vs *H*_1_ : *∃k, γ*_*k*_ *>* 0, we then propose to combine *q* + 1 single p-values corresponding to the *q* functional annotations and the original burden with ACAT^23^. Briefly, ACAT aggregates individual p-values and approximates the test statistic (and the subsequent p-value) based on a Cauchy distribution. So, for *q* + 1 tests in a region of interest, the ACAT statistic can be written as:

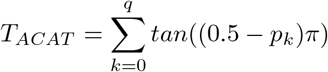

which follows approximately a Cauchy distribution under *H*_0_.

### 2.2 Bootstrap procedure using rare variant sharing patterns

We propose a weighted non-parametric bootstrap procedure in order to compute empirical p-values. Basically, genotypes were generated conditionally on the number of observed variants in a family, considering the rare variant sharing patterns occurring among family members. This procedure only requires the aggregated genotypes across affected individuals e.g.,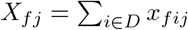 and the sharing pattern probabilities for a given family *f*, e.g., *P* (*G*_*fj*_). We apply the following procedure for estimating the null distribution of the test statistic:

- Sample aggregated genotypes for the *p*_*f*_ variants in family *f* across the *F* Families 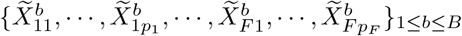 using the sharing pattern probabilities *P* (*G*_*f*_ *j*) obtained with RVS ^31^.
- Construct the test statistic 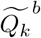.
- Compute empirical p-values for all *k*, 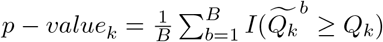
- ACAT-combined p-values are then obtained using empirical p-values instead of asymptotic p-values over the *q* + 1 functional annotations.

Because the *B* boostrap samples require only one set of rare variant sharing probabilities for all families, they only need to be computed once, hence increasing the computational performance, ensuring accurate estimation of p-values.

## 3 Numerical Simulations

We adopted the principle that CRHs are the annotations capturing best the causal variants, with simpler annotations capturing causal variants to a lesser extent. We thus selected a TAD in iPSC-derived neurons encompassing four CRHs showing different complexities (two genes-five enhancers (CRH1); two genes-two enhancers (CRH2); one gene-one enhancer (CRH3); one gene-four enhancers (CRH4)) to setup the simulation study. See Table S1 and ^30^ for more details. Genotypes were simulated based on observed variant sites and their corresponding MAF for the European population from the 1000 Genome Project database (phase 3). We extracted the 510 rare (MAF *≤* 1%) coding non-synonymous and within-enhancer non-coding single nucleotide variants from the TAD 800 kb (chr1:24100000-24970000). Using RarePedsim ^32^, we generated sequence data over the above 800 kb region for 270 affected subjects in the primary sample of 52 pedigrees ranging from small to extended (Figures 1 and S2) Families were simplified by removing inbreeding loops. For both Type I error rate and power evaluation, the dichotomous phenotype was assumed to follow a logistic model without covariates and with a population prevalence of 1%. Details on pedigree structures under the different scenarios were provided in Table S2. We focused on evaluating the ACAT-combined p-values. Importantly, to avoid large departures from the asymptotic distribution of RetroFun-RVS, we only considered functional annotations with a number of families greater than five. We also explored additional scenarios considering pedigrees of small to moderate size, families with a varying number of affected members and with presence of inbreeding. Details and results for these setups were provided in the Supplementary Numerical Simulations.

**Figure 1:**
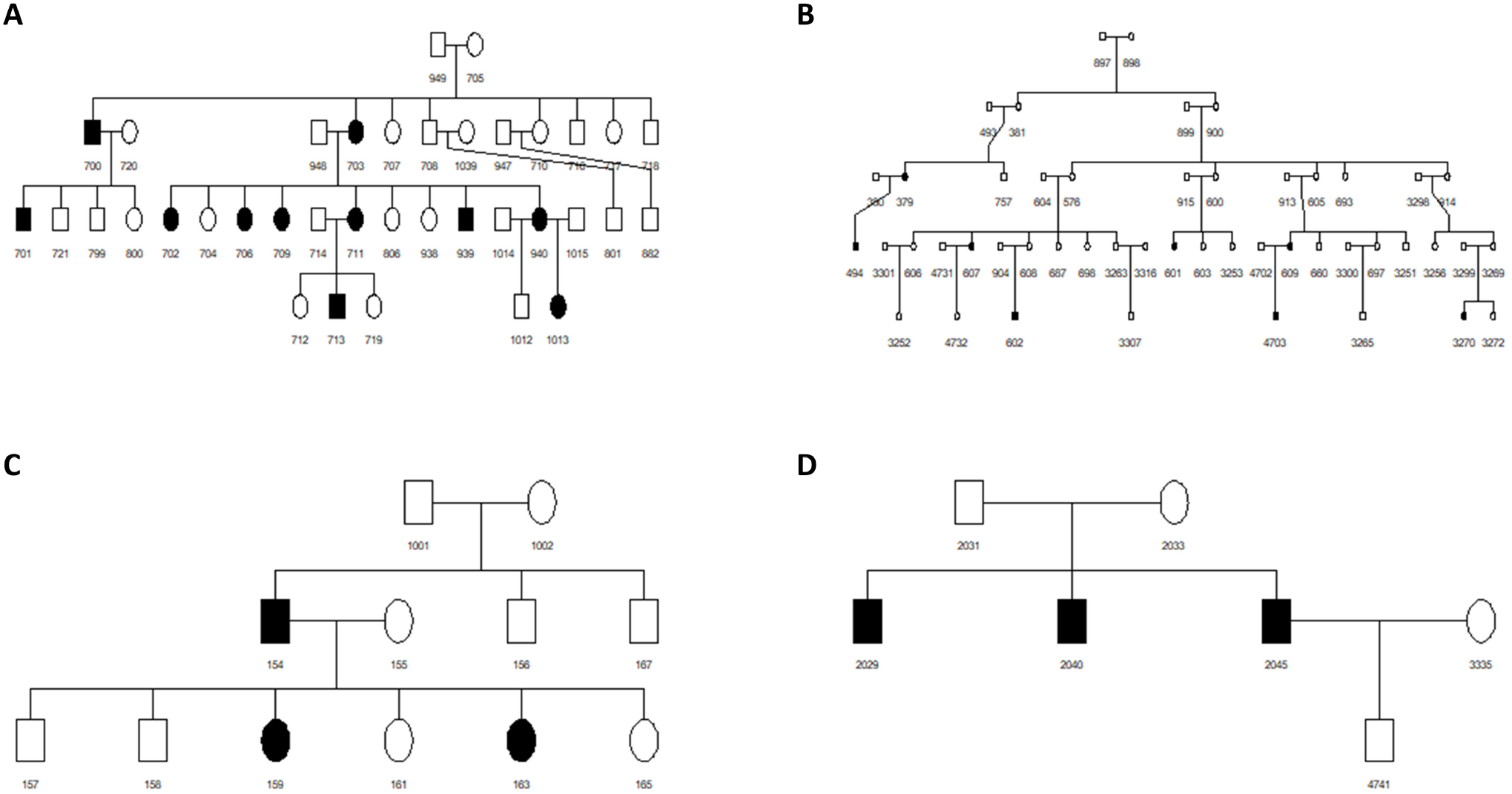
Example of pedigree structures considered in the simulation studies. Affected subjects are indicated by filled squares or circles

### 3.1 Type I Error Simulations

To determine whether the proposed framework preserves the desired Type I error rate, genotype data were generated unconditional on the affection status for family members. We specified a null effect for variants observed in families, i.e., odds-ratio (OR) = 1. Generating ten thousand replicates, we first examined the performance of RetroFun-RVS_*CRHs*_, which is RetroFun-RVS applied to CRHs and including variants over the entire TAD as global burden, with alternative definitions of regions to be included as functional scores: RetroFun-RVS_*Pairs*_, RetroFun-RVS_*Genes*_, and RetroFun-RVS_*Sliding−Window*_, for the method considering pairs of gene-enhancers, genes and a 10 Kb sliding window, respectively (Figure 2).

**Figure 2:**
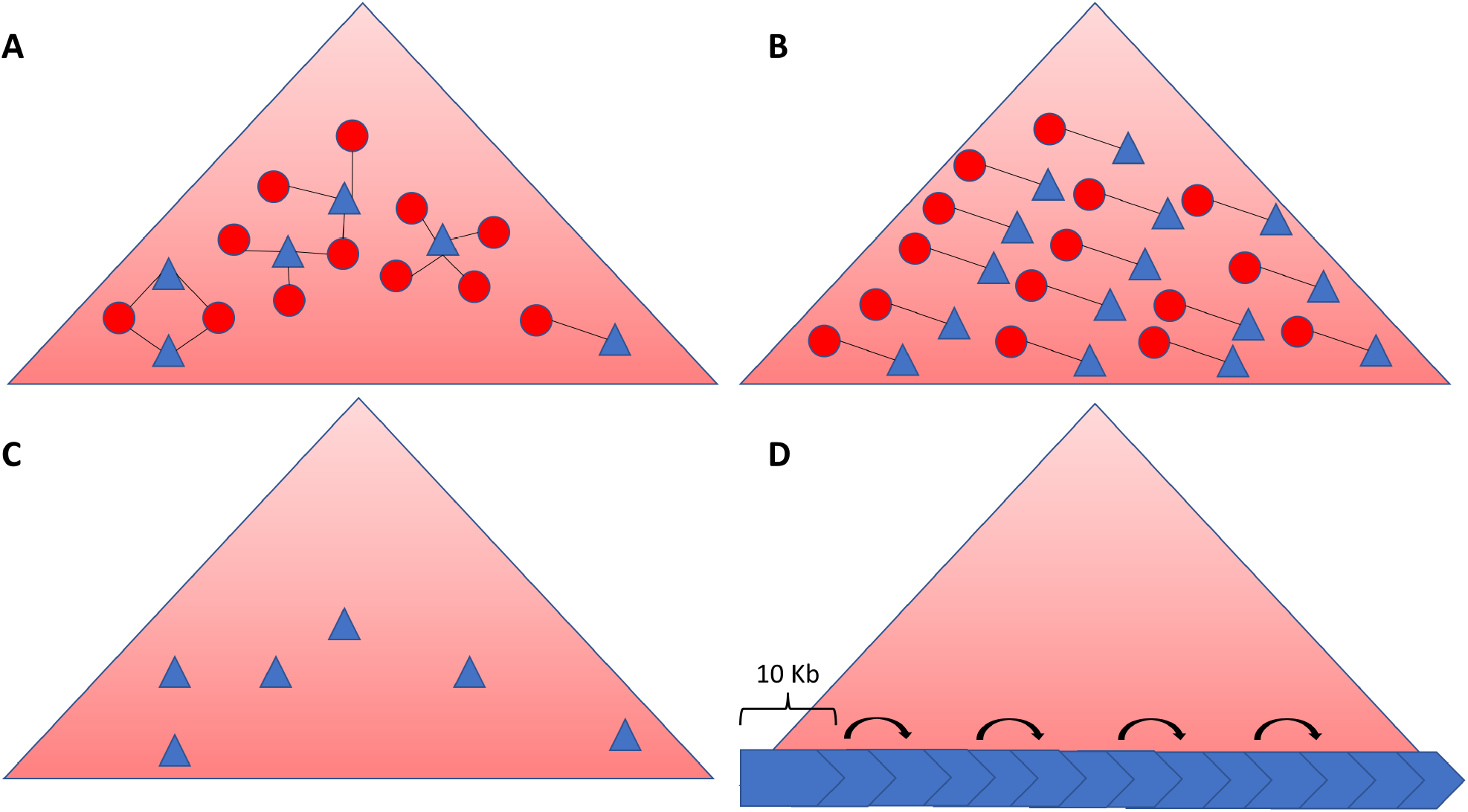
Overview of functional annotations considered in the simulation studies. For all 4 panels, big red triangles represent the selected TAD for the simulation studies, small blue triangles the genes (exons + promoters), and red circles the enhancers. (**A**) CRHs as functional annotations. (**B**) Pairs as functional annotations. CRHs are split with respect to each gene-enhancer pair. (**C**) Genes as functional annotations. (**D**) 10 Kb sliding windows as functional annotations.

### 3.2 Empirical Power Simulations

We set 2% of the variants over the entire region to be risk variants as suggested before ^28^, also performing simulations with 1% of risk variants as a sensitivity analysis. Genotypes were generated conditional on the affection status for each pedigree member assuming a multiplicative model with fixed variant effect, i.e., not depending on the MAF. Simulating one thousand replicates, we considered different scenarios where we varied the proportion of causal variants found in CRHs: 100%, 75% and 50% of causal variants (OR=5) were located within one CRH. The remaining variants being neutral (OR=1). This scenario is expected when variants are concentrated within elements functionally close. These three proportions correspond to the most advantageous scenario where all causal variants are within the same region and two mixed scenarios where signal is spread across the sequence of the region at different degrees. Our first evaluation assessed the gain of power by incorporating CRHs as functional annotations over the test including no scores (referred to as Burden Original). We also compared RetroFun-RVS_*CRHs*_ with others strategies to incorporate regions as functional annotations: RetroFun-RVS_*Pairs*_, RetroFunRVS_*Genes*_, and RetroFun-RVS_*Sliding−Window*_, for the method considering pairs of gene-enhancers, genes and a 10 Kb sliding window, respectively (Figure 2). Also, we assessed the performance in terms of power of our method compared to existing approaches namely, RVS ^15^ and RV-NPL ^17^ (Figure S3). Power was evaluated as the proportion of p-values less than *α* = 8.33 *×* 10^*−*6^, corresponding to the Bonferroni-adjusted 0.05 significance level when testing six thousand independent regions across the genome, corresponding to three thousand TADs (the average number of TADs found in our previous study across cell-types or tissues ^30^, while permitting the same number of additional domains of interest, i.e., outside TADs, to be tested. Results at lower proportion of risk variants and considering small pedigrees are also reported.

## 4 Illustration on the Eastern Quebec Schizophrenia and Bipolar Disorder Kindred Study

To illustrate the application of RetroFun-RVS to a whole-genome sequencing (WGS) study, we used data from the initial freeze of WGS on participants from the Eastern Quebec schizophrenia and bipolar disorder kindred study. Signed consent was obtained from all participants or from the parents for participants under 18 years of age for collection of all data analyzed here, under the supervision of the University-affiliated neuroscience and mental health ethics committee.

A description of genomic sequencing and data quality control can be found in the Supplementary Materials and Methods. For the present analysis we kept the 28 families with at least two relatives affected by the broad definition of schizophrenia, bipolar disorder and schizoaffective disorder in the Eastern Quebec Kindred Study ^33^. These 28 families included a total of 288 participants with WGS, including 175 who were affected. Inbreeding loops where two parents are first or second cousins were present in 6 families. All families of the Eastern Quebec kindred study were connected in a single genealogy with a mean completeness of 71% at the 10^*th*^ generation back using the BALSAC database (balsac.uqac.ca). Using that genealogy, we estimated to 0.0032 the mean kinship between the founders of the 28 families included in this study (the subjects who did not have parents in the 28 family structures before genealogy reconstruction). We used that value to apply the correction for cryptic relatedness to the RV sharing probabilities described by ^14^ in the computation of the expected value, variance and covariance of the score statistics *S*_*k*_(0) under the null hypothesis to obtain the asymptotic p-value of the tested variant sets. As a sensitivity analysis, we also analyzed the data using the standard approach described in subsection 2.1 in the simplified family structures without inbreeding loops from the primary sample used for the simulation study, replacing homozygous rare genotypes by heterozygous ones. The bootstrap procedure was applied to the variant sets yielding an aymptotic p-value below the significance level Bonferroni-corrected for the number of tests performed.

Our study focused on rare autosomal SNVs and short indels. We defined a rare variant as being absent or having a frequency < 0.01 in GnomAD non-Finnish European sample and in a sample of 1,756 controls from the founder Quebec population included in the CARTaGENE cohort (www.cartagene.qc.ca) ^34^.

We used as functional annotation the 1,633 CRHs defined by ^30^ in neurons derived from induced pluripotent stem cells. We included the 1237 CRHs covering at least one retained rare SNV or short indel, and either comprised in a single TAD (1042), overlapping two TADs (145) or outside any TAD (50). We applied ACAT to combine p-values of the burden test and CRH-specific tests in the 679 TADs with at least one rare SNV in a CRH entirely contained in the TAD and tested the other CRHs individually, for a total of 874 tests.

## 5 Results

### 5.1 Simulation of Type I Error Rate

The results show that, when we considered CRHs as functional annotations and accounted for variant dependence in the variance calculation, the Type I error rate was well-controlled when combining p-values using ACAT (Figure 3). However, we observed slight false positive inflations when RetroFun-RVS was applied with the independence variant structure or combining p-values using Fisher’s combined probability method (Figure S4A-S4B). Moreover, results for RetroFun-RVS_*CRHs*_ with no functional annotations and for each individual score show that the approach with covariance terms is either well calibrated or slightly conservative (Figure S4D to S4F). In addition, the method shows moderate Type I error rate inflation when applied to small to moderate family structures, increasing when assuming variant independence (Figure S5). Further investigations have shown that Type I error depends on the structure considered (Figure S6). When investigating scenarios in presence of inbreeding, we observed that RetroFun-RVS_*CRHs*_ considering homozygous configurations slightly reduces Type I error inflation in presence of a modest number of inbred families, compared to results where consanguinity is left untreated (Figure S7A), consistent with the improvement in Type I error control achieved by the dependence correction. In contrast to our “only-inbred” scenario, where high level of false positives are observed even when considering homozygous configurations (Figure S7B).

**Figure 3:**
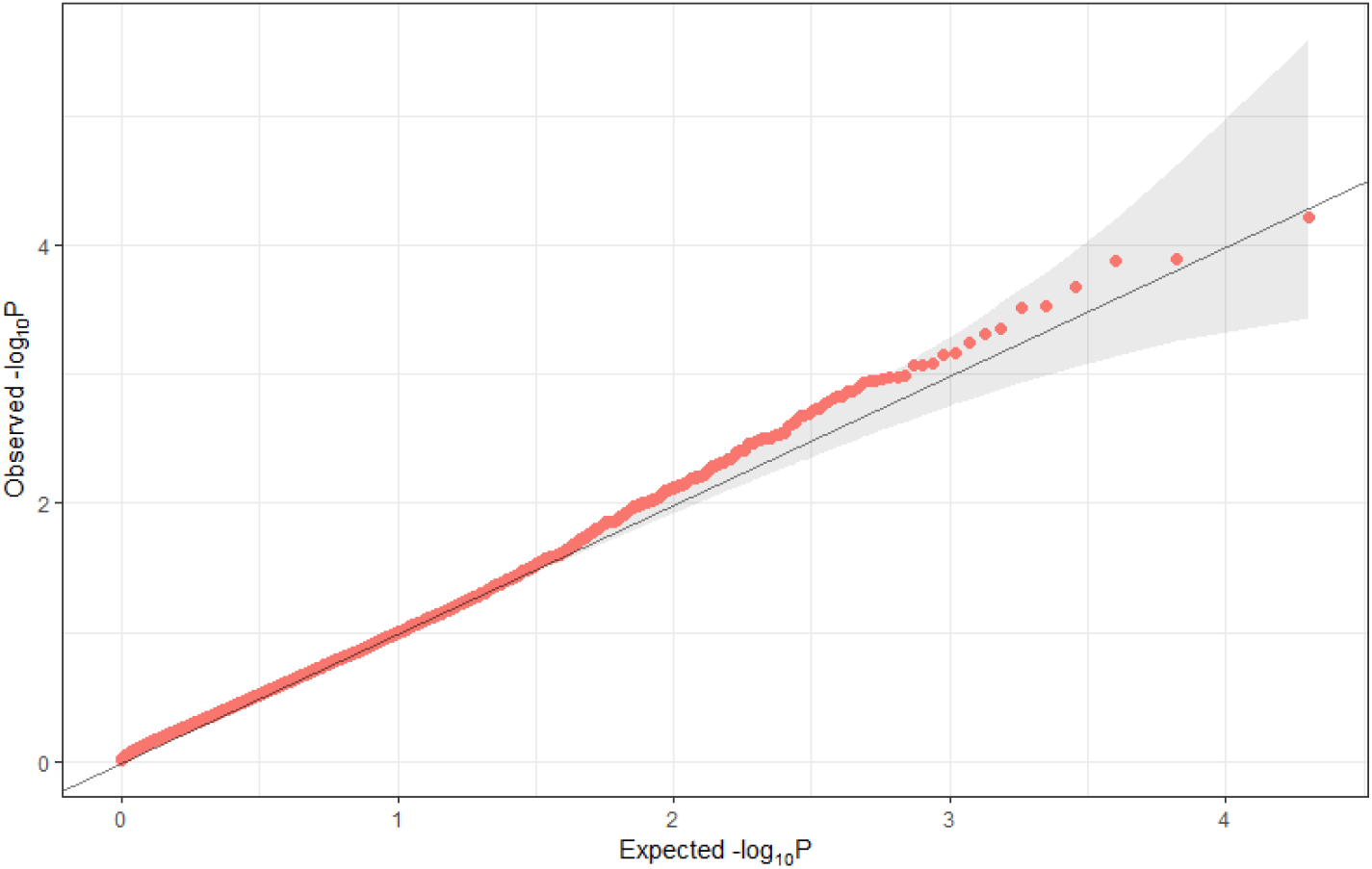
Quantile-Quantile plot of ACAT-Combined P-values for RetroFun-RVS_*CRHs*_ considering variant dependence. We omitted CRHs with a number of families less than 5 ensuring a proper asymptotic behavior.

Turning now to pairs and genes as functional annotations, we observed moderate inflation of the Type I error rate in extended pedigrees, even when considering variant dependence, while for 10Kb sliding windows the Type I error rate inflation was more severe (Figure S8). We attempted to discard 10 kb windows with few variants, and observed that Type I error control was achieved on windows encompassing 30 variants or more but few windows met this requirement (results not shown). Moreover, the bootstrap procedure applied to RetroFun-RVS_*Pairs*_, RetroFun-RVS_*Genes*_ and, RetroFun-RVS_*Sliding−Window*_ to compute p-values empirically provides Type I error rate control, although being conservative, particularly for functional annotations encompassing few variants (Figure S9). To summarize, the results show that RetroFun-RVS with asymptotic p-values is a valid approach when CRHs or a large region are considered in extended pedigrees, despite being inflated to various degrees for others strategies or certain family structures. Bootstrap p-values can be computed in these instances to control the Type I error rate.

### 5.2 Power and Scalability Comparison Considering Different Strategies to build Functional Annotations

In the first set of power evaluations, we assessed power under different scenarios of causal variant distributions. Firstly, we compared RetroFun-RVS integrating CRHs with the same method incorporating no functional annotation. Consequently, when 100% and 75% of causal variants were within one CRH, our method RetroFun-RVS_*CRHs*_ performed better than the original burden test showing gains of 10% and 9%, while at 50% causal the power remains comparable (Figure 4A). Also, considering only pedigrees of small to moderate size, we observed that, even if both RetroFun-RVS_*CRHs*_ and the original burden test without annotation exhibit lower power, the gain for RetroFun-RVS_*CRHs*_ becomes higher as the percentage of causal variant within the CRH of interest increases (Figure S10). Congruent results were obtained when a lower proportion of causal variants was considered, showing a minimal power gain of 10% and a maximal increase of 125% (Figure S11). Therefore, our findings suggest that substantial power gain can be achieved when CRHs are predictive for the effect of variants on the trait, RetroFun-RVS_*CRHs*_ showing robustness when signal is spread across several CRHs. Then, we compared RetroFun-RVS_*CRHs*_ to other strategies to integrate regions as functional annotations, namely RetroFun-RVS_*pairs*_, and RetroFun-RVS_*genes*_. Our results show that integrating CRHs as functional annotations is a more powerful strategy compared to the other strategies considered (Figure 4B). The power of RetroFun-RVS considering sliding windows as functional annotations comparable to RetroFun-RVS_*CRHs*_ (Figure S12) is likely explained by inflated Type I error rate. Globally our results follow the same pattern when decreasing the proportion of causal variants (Figure S13). In summary, RetroFun-RVS_*CRHs*_ exhibits power gains when CRHs show high or modest percentages of causal variants. Also, the method is robust and powerful under the different scenarios that we considered, that are, inclusion of weakly predictive CRHs, small percentages of risk variants, and presence of small families.

**Figure 4:**
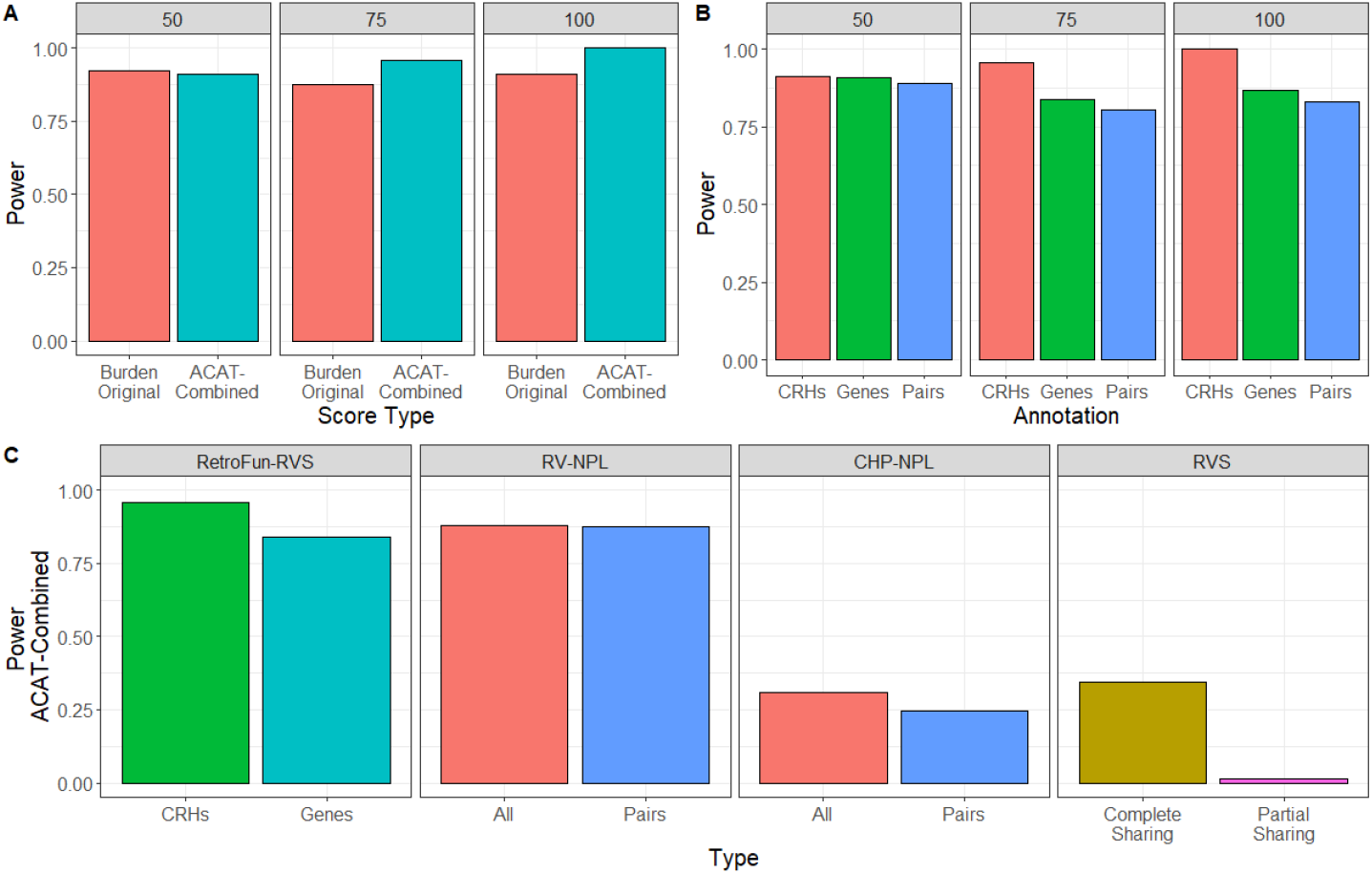
Power evaluation of RetroFun-RVS under different scenarios for 2% risk variants. (**A**) Power at different proportions of risk variants within the CRH, between RetroFun-RVS_*CRHs*_ with no functional annotation (Burden Original) and RetroFun-RVS_*CRHs*_ including the four CRHs (ACAT-Combined). Power was evaluated on 1,000 replicates. (**B**) Power at different proportions of risk variants within the CRH between RetroFun-RVS_*CRHs*_ (CRHs), RetroFun-RVS_*Pairs*_ (G-E Pairs), RetroFun-RVS_*Genes*_ (Genes), and RetroFun-RVS_*Sliding−Window*_ (Sliding). Functional annotations with fewer than five families were removed from the analysis for ensuring a proper asymptotic behavior. Given the Type I error inflation observed for RetroFun-RVS_*Sliding−Window*_, this approach was excluded from the power comparison. Power was evaluated on 1,000 replicates. (**C**) Power at 75% risk variants within one CRH between RetroFun-RVS_*CRHs*_ and other affected-only competing methods. Here we included RetroFun-RV_*genes*_ to mimic CHP-NPL procedure. Power for RetroFun-RVS_*CRHs*_ and RetroFun-RVS_*Genes*_ was evaluated on 1,000 replicates, while for RV-NPL and RVS we generated 200 replicates.

### 5.3 Power Comparison with Others Affected-Only Methods

In the second set of power evaluations, we compared RetroFun-RVS_*CRHs*_ with other affected-only methods, namely RVS ^15^ and RV-NPL ^17^. Thus, to proceed to fair comparisons between methods, we adapted RVS and RV-NPL to take CRHs into account (See Supplementary Numerical Simulations). With 2% risk variants, when we considered 75% of causal variants located within one CRHs, we observed that RetroFun-RVS reaches greater power compared to competing methods 4C), exhibiting significantly shorter computing times (Table 1). At lower proportions of risk variants, the new method remains more powerful compared to RV-CHP or RVS, and equivalent to RV-NPL (Figure S14).

**Table 1:**
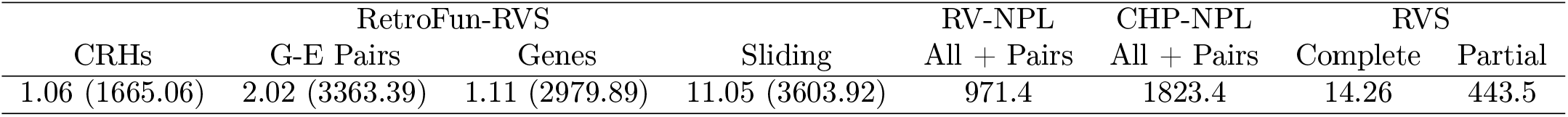
Running times (in seconds) for analyzing rare variants in the TAD, in one simulated replicate, using a single 2.10GHz processor. For RetroFun-RVS, we also provided average running times for computing empirical p-values based on 10,000 samples (in parenthesis). For RV-NPL empirical p-values were obtained based on 1 million permutations.

## 6 Illustration on the Eastern Quebec Schizophrenia and Bipolar Disorder Kindred Study

No ACAT-combined-over-TAD or single CRH p-value reached the significance level *α* = 0.05*/*874 = 5.7 *×* 10^*−*5^ corresponding to a Bonferroni correction for the number of tests performed, after recomputing with the bootstrap the asymptotic p-values below that level. As an illustration, we provide details of the CRH with a bootstrap p-value = 0.00016 (asymptotic p = 0.000077) to illustrate patterns of sharing that can be captured by RetroFun-RVS. The original Burden p-value is 0.18, thus if variants in the CRH are true susceptibility variants, this result would be aligned with our simulation studies in which the unified test was more powerful with predictive functional annotations. This CRH between positions 43998889 and 44492786 on chromosome 7 encompasses 11 genes (*PGAM2, POLM, AEBP1, DBNL, POLD2, RASA4CP, YKT6, CAMK2B, SPDYE1, NUDCD3, POLR2J4*) and 19 enhancers. Importantly, on the eight variants seen in at least one affected subject (in a total of eleven families), four were located either in an intergenic or a genic enhancer, impacting between one and ten genes simultaneously. These enhancers located up to 343 Kb distance apart from their target genes (average 91Kb). This result suggests that strategies linking non-coding variants to the nearest gene will fall short in identifying the putative causal gene. We illustrated this result in Figure S15. Figure S16 illustrates the family with the rare SNV shared by the most affected subjects, including one who shares the rare allele with other affected family members through unknown relations accounted for by the correction for cryptic relatedness based on the kinship among founders.

## 7 Discussion

Most of rare genetic variations are located within non-coding regions, making the underlying biological mechanisms through which they impact disease risk difficult to interpret. Over the past few years, efforts were not only made in annotating the genome but also integrating these annotations into statistical frameworks ^21,22^. Although such methods have already been developed for unrelated subjects such as case-control samples, to our knowledge, no approach to date has been proposed to integrate functional annotations within family-based designs. In this paper we have presented RetroFun-RVS, a retrospective burden test, integrating functional annotations considering only affected individuals within families. We have shown that binary annotations corresponding to disjoint regions with regulatory impacts, such as CRHs, provide power gains when such regions concentrate causal variants, outperforming other strategies or competing methods (Figure 4), while well controlling the Type I error rate in samples of families of various size and structure (Figure 3). Since regulatory mechanisms are highly tissue- or context-dependent it can be challenging to have the right tissue for the right trait, and misspecifying the model is likely in practice. Thus, integrating the original burden test, corresponding to aggregating all variants across a region, in RetroFun-RVS makes it robust, showing stable power when functional annotations poorly predict the trait. Finally, by computing p-values asymptotically, RetroFun-RVS is computationally faster than competing methods, which often require permutation-based approaches or exact probability computations to sharply control the Type I error rate.

The main rationale behind RetroFun-RVS is that risk variants are enriched among affected individuals compared to the expected variant count based on their relationships. Hence, one critical feature of our method is to aggregate genotypes while measuring rare variant sharing among affected family members to compute the test statistic. To implement an affected-only analysis, where individuals are selected based on their disease status, we have adopted a retrospective approach, considering genotypes as random, while conditioning on phenotypes ^18^. Also, since genotype probabilities do not depend on MAF under the assumption that the variant frequency tends to 0, RetroFun-RVS necessitates only familial information to compute these probabilities, in order to derive the score statistic and its variance (See Material and Methods). This aspect is central, since the variance terms need to be computed only once for the entire set of families, which is computationally efficient even in presence of large pedigrees. Our rare variant assumption however implies that genotypes homozygous for the rare allele are impossible in the absence of inbreeding. Data simulated for Type I error and power assessments did contain the small number of homozygous rare genotypes expected for variants with MAF = 1%. Conversion to heterozygous genotypes did not increase Type I error rate compared to removing the variants with homozygous rare genotypes, so we only showed results with the conversion to heterozygous genotypes. Also, we observed that RetroFun-RVS controls the Type I error rate well in the presence of a small to modest number of inbred families (Figure S7A). Thus, if in practice cryptic relatedness or inbreeding are expected for only a small proportion of families, we suggest to use RetroFun-RVS without considering homozygous configurations. Indeed, depending on the inbreeding configuration, computational times may be extremely long, limiting applications for large families. Moreover, our simulation studies have demonstrated that in the presence of a high proportion of inbred families or a high level of inbreeding, RetroFun-RVS may suffer from severe inflation without allowing homozygous configurations, and some inflation remains even when handling homozygosity (Figure S7B). In an intermediate scenario such as the application to the Eastern Quebec Kindred Study with widespread cryptic relatedness and some inbreeding, considering homozygous configurations may provide a gain in power and is recommended.

In addition to being computationally effective, RetroFun-RVS is more powerful than other affected-only competing methods, under certain scenarios (Figure 4C, Figure S13). For example, compared to RVS, on which RetroFun-RVS is built upon, but which can only analyze between one and five rare variants simultaneously in the pedigree sample used in the simulation study, we reached greater power by testing tens of variants together in annotated regions, or even hundreds of variants in the absence of annotations. It is noteworthy that the simulated variant ORs did not depend on the variant MAF due to limitations of the simulation software. The MAF-dependent variant weighting scheme of RetroFun-RVS was thus misspecified in the power evaluation. Greater power gains of RetroFun-RVS over the competing methods ignoring variant MAF could have been achieved had the variant ORs be inversely related to MAF.

Although the score test was well-calibrated and powerful in our primary sample covering a large size range from small families to extended pedigrees, we have detected modest Type I error rate inflation with another sample of small to moderate family structures (Figure S5). Additional investigations have shown that RetroFun-RVS controls the false positive rate accurately under certain family structures, while providing slightly conservative or inflated quantile-quantile plots for other structures (Figure S6). Since we did not observe clear associations between the number of affected individuals and false positive rates, we argue that the inflation observed is more a question of family structure than family size. We argue that RetroFun-RVS controls the Type I error rate adequately with typical family samples consisting in combinations of small to extended pedigrees. On a related note, some analyses have shown that Type I error rate or power are highly dependent on the number of variants present in the region of interest. Indeed, we have observed that when large numbers of variants are considered, RetroFun-RVS might provide conservative results involving some power loss (Figure S4C), while a small number of variants tends to offer inflated Type I error rate (Figure S8). Complementary analyses are needed to inspect the empirical relationship between size of region and performance. We recommend in practice to use the dependence-adjusted model. Bootstrap procedures (Figure S9) might be considered to sharply control Type I error rate when unsure of Type I error control due to pedigree structures or for small numbers of variants at the expense of longer computing time. However, since only small p-values are relevant, application of the bootstrap can be limited to the hits obtained from the asymptotic p-value computation, mitigating the computational requirements. Interestingly, the non-parametric bootstrap procedure offers faster running times for generating 10,000 samples when considering CRHs as functional annotations, ranging from the single to double, depending on the type of annotations considered (Table 1).

Moreover, RetroFun-RVS in its current form is restricted to binary phenotypes and does not allow the integration of individual-level covariates, such as sex, age or genetic principal components. Hence, future work is needed to extend the framework to cases selected by extreme values of continuous phenotypes and to include covariates.

We argue that the performance of the proposed method is strongly dependent to the availability of the relevant tissue for the studied disease. Indeed, regulatory mechanisms operate in a tissue- or cell-type-specific manner. Our framework, by allowing the incorporation of several functional annotations from diverse tissues or cell-types without loss of power, is useful to highlight the underlying biological mechanisms involved in the trait. This aspect is central from a fine-mapping perspective, thus RetroFun-RVS will be an important tool to pinpoint causal variants located within non-coding regions, which could have been missed so far.

## 8 Data and Code Availability

Cis-Regulatory Hubs and Topologically associated domains used in this paper are available on https://github.com/lmangnier/CRHs. Variant data were available from the 1000 Genome project: https://www.internationalgenome.org/data-portal/data-collection/phase-3. The data of the Eastern Quebec SZ and BD kindred study are available on request from the corresponding author. We have implemented RetroFun-RVS in a R package, available on GitHub (https://github.com/lmangnier/RetroFun-RVS). The code for the simulation study is at https://github.com/lmangnier/Simulation_RL and the code for the processing and analysis of the Eastern Quebec schizophrenia and bipolar disorder kindred study whole genome sequence data is at https://github.com/abureau/RV_in_SZ_BD_kindreds.

## 9 Conflict of Interest

The authors declare that they have no conflict of interest.

## Notes

https://github.com/lmangnier/Simulation_RetroFun-RVS

https://github.com/lmangnier/RetroFun-RVS

### Competing Interest Statement

The authors have declared no competing interest.

### Summary of Updates

Author affiliations updated Supplemental files updated Clarify Introduction Clarify Discussion Method section updated to provide right-tail test, clarify underlying assumptions of the model and bootstrap sub-section Result section updated to provide: - Additional simulation scenarios for evaluating Type-I error rate across different structures of families and in presence of inbreeding - Applications to whole genome sequencing data in extended pedigrees for schizophrenia and bipolar disorders Figure 3 updated Figure 4 updated

