## Supplementary Material for "RetroFun-RVS: a retrospective family-based framework for rare variant analysis incorporating functional annotations"

### 1 Material and Methods

#### 1.1 Allowing homozygous genotypes for rare alleles in case of inbreeding or cryptic relatedness

The main requirement to allow homozygous genotypes for rare alleles in the computation of the expected value, variance and covariance of the RetroFun-RVS score statistic is to compute probabilities for all $3^{n_f} - 1$ genotype configurations $G$ conditional on observing at least one copy of the rare allele in family $f$ (using the option `distinguishHomo` of the `RVsharing` function of the `RVS` package) instead of restricting the computation to the $2^{n_f} - 1$ configurations with only heterozygous and homozygous common genotypes. Homozygous genotypes for rare alleles have positive probability in inbred individuals under the assumption that only one copy of the minor allele was introduced once by a family founder, and have positive probability in all individuals when allowing rare variants to be introduced twice in the family by two distinct founders with a probability depending on the mean kinship in the population from which the founders are drawn, an approximate correction of the RV sharing probabilities in case of cryptic relatedness among founders, available in the `RVsharing` function. Variants with homozygous genotypes for their rare allele are included in the RetroFun-RVS score statistic when homozygous rare genotype probabilities are computed by specifying an option of the `RetroFunRVS` function. It is recommended to use the cryptic relatedness correction in presence of inbreeding as it allows positive probabilities of homozygous rare genotypes for all family members, and as relatedness among founders is likely in populations where inbreeding is prevalent.

#### 1.2 Risk variants dominate protective variants in an affected-only design

To assess whether in an affected-only design, the contribution to the score statistic of a risk variant dominates the contribution of a protective variant with equal opposite effect, we considered two different scenarios corresponding to two different family structures where (1) one pair of first cousins are affected and (2) one pair of second cousins are affected.

Here we perform the calculations for one variant and for a pair of first cousins. Let's define the sharing probability under the null $p_0 = 1/15$. Hence, the expected value under the null for this family configuration is given by: $\lambda_0 = 2 * p_0 + 1 * (1 - p_0) = 1.06$. As an example, assuming a relative risk (RR) of 50 the expected statistic value under the alternative for a risk variant is $(1 - \lambda_0) * P(G_{.j} = 1) + (2 - \lambda_0) * P(G_{.j} = 2)$, where $P(G_{.j} = 2) = 1/(1 - p_0)/(50 * p_0 + 1)$, while for a protective variant $P(G_{.j} = 2) = 1/(1 - p_0)/(1/50 * p_0 + 1)$, which gives 0.71 and -0.06 respectively. Thus, we observed that for a risk variant the statistic increases with effect size while for protective variant, the statistic remains stable. We can conclude that risk variants contribute more to the score statistic than protective variants (See Figure S1 below).

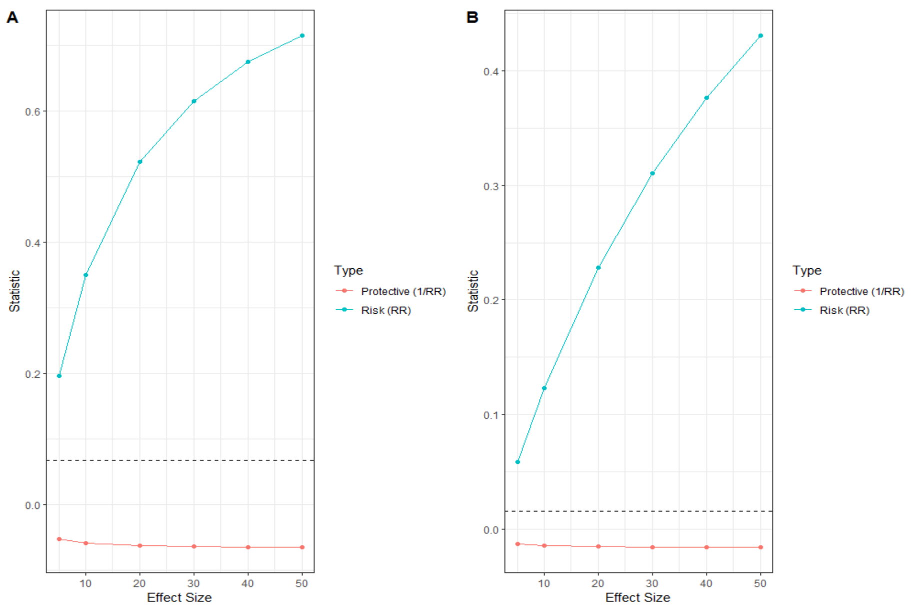

Figure S 1: Expected contribution of a family to the score statistic value for different effect sizes considering either 1 risk variant (RR) or 1 protective variant (1/RR). (**A**) Pair of affected first cousins, (**B**) Pair of affected second cousins. The sharing probabilities under the null, assuming variant frequency tending to 0, are 1/15 and 1/63, respectively, and indicated by the dashed horizontal lines

#### 1.3 Score Variance

The score variance is given by:

$$Var[S_k(0)] = Var[\sum_j w_j Z_{jk} \sum_i X_{fij}]$$

$$= \sum_f \sum_j w_j^2 Z_{jk}^2 Var[X_{f.j}] + \sum_{j' \neq j} w_j w_j' Z_{jk} Z_{j'k} Cov[X_{f.j}, X_{f.j'}]$$

where $X$ is the random variable corresponding to the minor allele count $x$ and $X_{f.j} = \sum_i X_{fij}$.

Developing variance and covariance components respectively we obtained:

$$Var[X_{f.1}] = \sum_x x^2 P(X_{f.1} = x) - \left( \sum_x x P(X_{f.1} = x) \right)^2$$

and

$$Cov[X_{f.1}, X_{f.2}] = \sum_{x_1} \sum_{x_2} x_1 x_2 P(X_{f.1} = x_1; X_{f.2} = x_2) - \left( \sum_x x P(X_{f.1} = x) \right)^2$$

Since these components are identical for all variants and variant pairs, they need to be computed only once for all variants.

Three possible cases can arise for the covariance term:

- Variants are in perfect linkage disequilibrium and we have $Cov[X_{f.1}, X_{f.2}] = Var[X_{f.1}]$. This is treated in practice by keeping only one of the two variants since their genotypes are identical. The default implementation keeps the variant with the highest annotation value (which assumes that the values of different annotations are comparable; it is the case with binary annotations such as CRHs).

- Variant independence can be assumed and $Cov[X_{f.1}, X_{f.2}] = 0$.

- General case of imperfect linkage disequilibrium. The term $P(X_{f.1}; X_{f.2})$ would be difficult to compute since we condition on $X_{f.j} \geq 1, j \in V_f$ where $V_f$ is the subset of variants in family $f$. Instead, we set

$$Cov[X_{f.1}, X_{f.2}] = \sum_{x_1} \sum_{x_2} x_1 x_2 \frac{\min(P(X_{f.1} = x_1), P(X_{f.2} = x_2))}{\sum_{u,v} \min(P(X_{f.1} = u), P(X_{f.2} = v))}$$
$$-(\sum_{x} xP(X_{f.1} = x))^2$$

  to obtain an approximation of the variance of the score $S_k(0)$.

#### 2 Numerical Simulations

| CRH1 | CRH2 | CRH3 | CRH4 | Out |
|---|---|---|---|---|
| 158 | 65 | 2 | 41 | 244 |

Table S 1: Number of variants located within each CRH and outside.

#### 2.1 Pedigree Structures

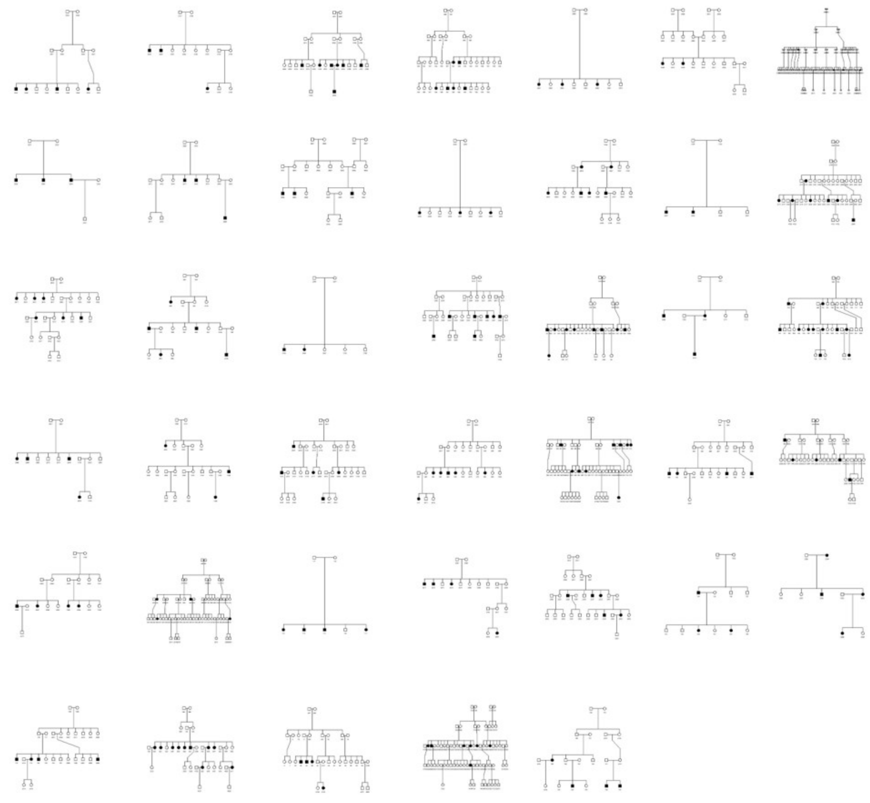

Figure S 2: Pedigree structures for all the 52 families considered in the simulation studies. Affected subjects are indicated by filled squares or circles. 12 pedigree structures are repeated (see Table S2)

#### 2.2 Additional simulation scenarios

As mentioned in the main manuscript, we generated additional simulation scenarios for evaluating the performance of RetroFun-RVS with respect to the control of the Type I error rate under different pedigree structures. Pedigree structures for main and secondary simulation settings were summarized in Table S2.

- Firstly, we considered one additional scenario with only pedigrees of small to moderate size, restricting our attention to the 17 families with fewer than 10 individuals. However, we repeated these families several times to reach the same total number of affected subjects as in the primary sample.

- Since we observed slight inflation of the Type I error rate under our scenario with small to moderate pedigrees, we produced six additional simulation settings, restricting our attention to families with a number of affected members from two to seven. We repeated families several times to achieve roughly the same number of affected subjects as in the primary sample.

- In practice pedigrees can exhibit either inbreeding or cryptic relatedness leading to the violation of the 'one founder one copy' assumption of RVS. Thus, for evaluating the performance of RetroFun-RVS to control the Type I error rate in the presence of homozyguous configurations, we produced two additional scenarios with (1) inbred and non-inbred pedigrees and (2) only inbred families. Albeit rarely encountered in practice, this latter scenario served us as an extreme case for assessing the control of the Type I error rate. RetroFun-RVS was evaluated with and without correction for homozyguous configurations under these two settings.

#### 2.3 Adaptation of RVS and RV-NPL including CRHs

Given the computational complexity of RVS is exponential in the number of variants tested together, we had to restrict the RVS tests to short windows. In order to adapt RVS to take CRHs into account, variants were sorted such that all variants belonging to the first CRHs encountered in the region are in consecutive order, followed by the variants in the second CRH and so on until the last CRH (Figure S3). Then, at the end, variants outside CRHs are listed. Windows were defined as sets of consecutive variants in that order, such that windows can span multiple elements of the same CRH (e.g. elements 1 and 2 of the blue CRH on (Figure S3). Computation of RVS tests over windows of 5 variants were attempted. When memory was insufficient, the rightmost variant was removed and the computation reattempted. Variants were removed until computation was successful. The window reduction was only needed for the partial sharing RVS test; the complete sharing RVS test computation was always successful on windows of 5 variants.

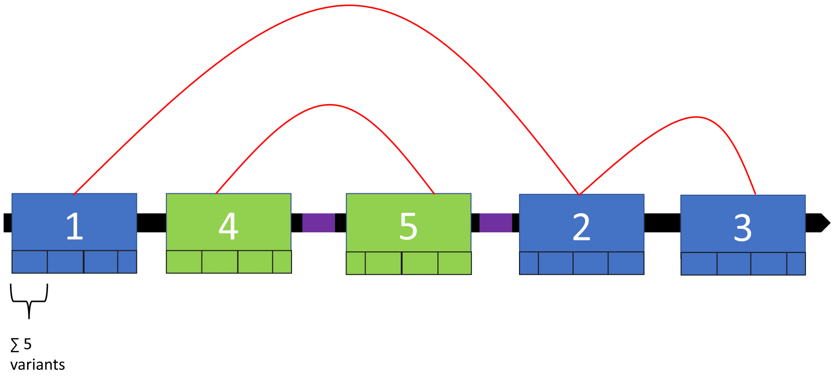

Figure S 3: Adaptation of RVS including CRHs. The blue and green regions are two distinct CRHs and the black regions are outside any CRH. Purple rectangles correspond to windows outside CRHs.

We also adapted RV-NPL to integrate CRHs. Variants belonging to each CRH were specified through the option *include-vars* using **rvnpl** *collapse* when generating results from both RV-NPL and CHP-NPL. Moreover, to obtain an unified p-value, we applied ACAT on p-values from each CRH for RV-NPL and CHP-NPL and p-values from each window for the RVS tests.

#### 2.4 Genomic sequencing and data quality control

Freshly extracted DNA from stored frozen blood leukocytes was used for WGS. After TruSeq DNA PCR-free library preparation, sequencing was performed with the short read NovaSeq technology from Illumina using S4 flowcells version 1.5. Each individual sample was sequenced with a target 30X coverage, which was achieved for 64.2% of the samples, while all samples had coverage ¿ 25X. We used the Illumina DRAGEN Germline pipeline to align reads to H. sapiens hg38 reference genome and then performed a joint calling of variants on the 349 participants with sufficient coverage from 40 families with the DRAGEN Popgen pipeline (v.3.10.4).

Variants with missingness $> 0.05$ and those with an undefined alternative allele were removed. Mendelian inconsistencies were detected and resolved by removing the genotypes of the involved individuals. Monomorphic variants were then removed. The genomic mask for hg38 derived from the 1000 Genomes Project phase 3 data [1] was then applied to keep only the variants in mappable regions.

We computed on each autosomal chromosome a scan statistic [2] to detect regions where variants absent in the CARTaGENE cohort form clusters, and removed such variants located in clusters detected with $p < 0.05/22$ to control for the number of chromosomes analyzed. In addition, we removed variants seen in members from more than 20% of the families included in the analyses ($\geq 6/28$). After application of all these filters, there remained 1,913,282 rare SNVs and short indels.

### 3 Results

#### 3.1 Type I Error Rate

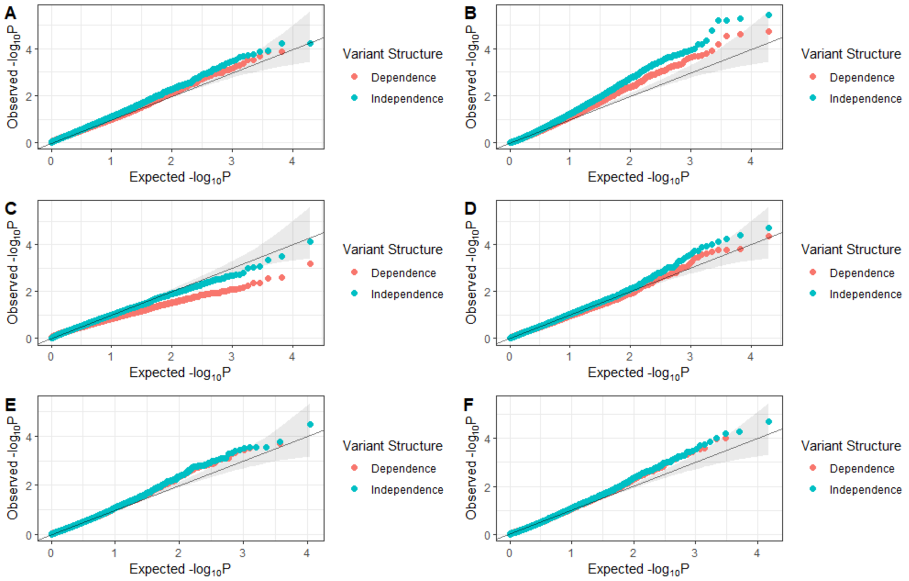

Figure S 4: (**A**) Quantile-Quantile plot of ACAT-Combined p-values for RetroFun-RVS$_{CRHs}$ considering variant dependence and independence. (**B**) Quantile-Quantile plot of Fisher-Combined p-values for RetroFun-RVS$_{CRHs}$ considering variant dependence and independence. (**C**) Quantile-Quantile plot for RetroFun-RVS$_{CRHs}$ considering the original Burden test under variant dependence and independence. (**D**) Quantile-Quantile plot for RetroFun-RVS$_{CRHs}$ considering the first CRH test under variant dependence and independence. (**E**) Quantile-Quantile plot for RetroFun-RVS$_{CRHs}$ considering the second CRH under variant dependence and independence. (**F**) Quantile-Quantile plot for RetroFun-RVS$_{CRHs}$ considering the fifth functional annotation under variant dependence and independence. As mentioned in the main manuscript, functional annotations with fewer than five families were removed from the analysis for ensuring a proper asymptotic behavior. Because only a few replicates had p-values for CRH 3, we omitted it in the analysis. Overlapping points (where p-values from both methods are identical) appear as turquoise.

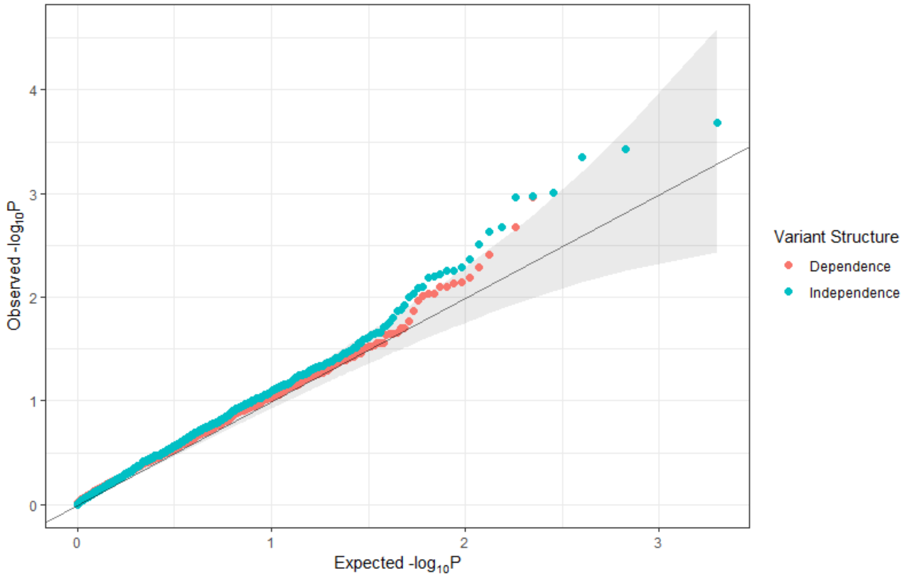

Figure S 5: Quantile-Quantile plots of ACAT-Combined p-values for RetroFun-RVS$_{CRHs}$ considering only pedigrees of small to moderate size under variant dependence and independence structure. As mentioned in the main manuscript, functional annotations with fewer than five families were removed from the analysis for ensuring a proper asymptotic behavior. 1,000 replicates were generated for this scenario. Overlapping points (where p-values from both methods are identical) appear as turquoise.

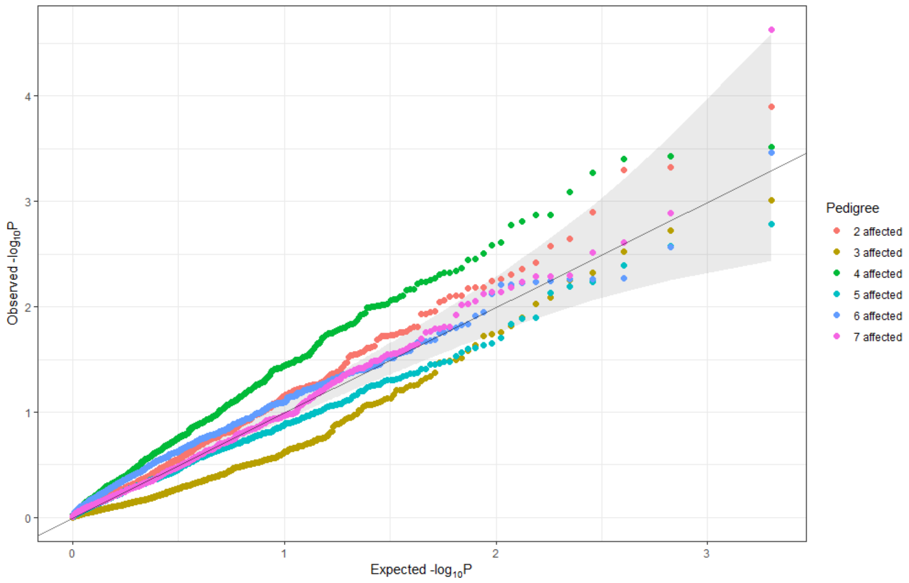

Figure S 6: Quantile-Quantile plots of ACAT-Combined p-values considering different pedigree structures under variant dependence. As mentioned in the main manuscript, functional annotations with fewer than five families were removed from the analysis for ensuring a proper asymptotic behavior. 1,000 replicates were generated for this scenario.

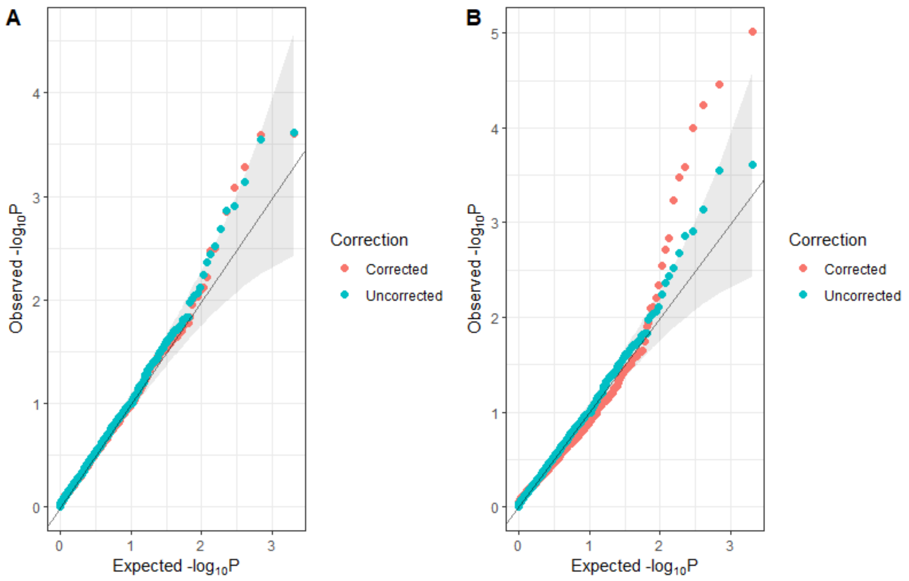

Figure S 7: Quantile-Quantile plots of ACAT-Combined p-values considering variant dependence where (**A**) inbred and non-inbred families were analyzed and where (**B**) only inbred families were analyzed. Under these scenarios we allowed for homozygous genotypes. As mentioned in the main manuscript, functional annotations with less than five families were removed from the analysis for ensuring a proper asymptotic behavior. 1,000 replicates were generated for this scenario. Overlapping points (where p-values from both methods are identical) appear as turquoise.

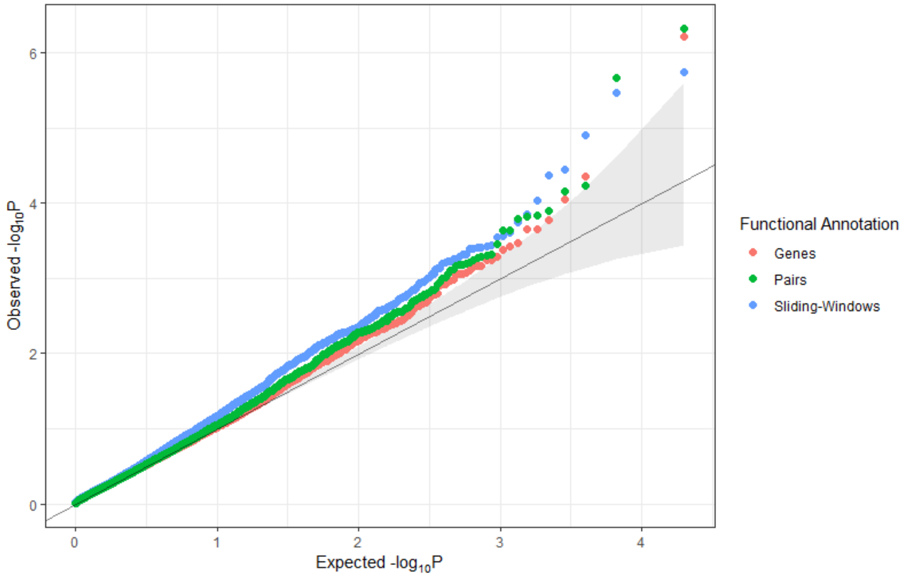

Figure S 8: Quantile-Quantile plots of ACAT-Combined p-values considering variant dependence for: RetroFun-RVS$_{Pairs}$, RetroFun-RVS$_{Genes}$, and RetroFun-RVS$_{Sliding-Windows}$. As mentioned in the main manuscript, functional annotations with fewer than five families were removed from the analysis for ensuring a proper asymptotic behavior. 1,000 replicates were generated for this scenario. Overlapping points (where p-values from both methods are identical) appear as turquoise.

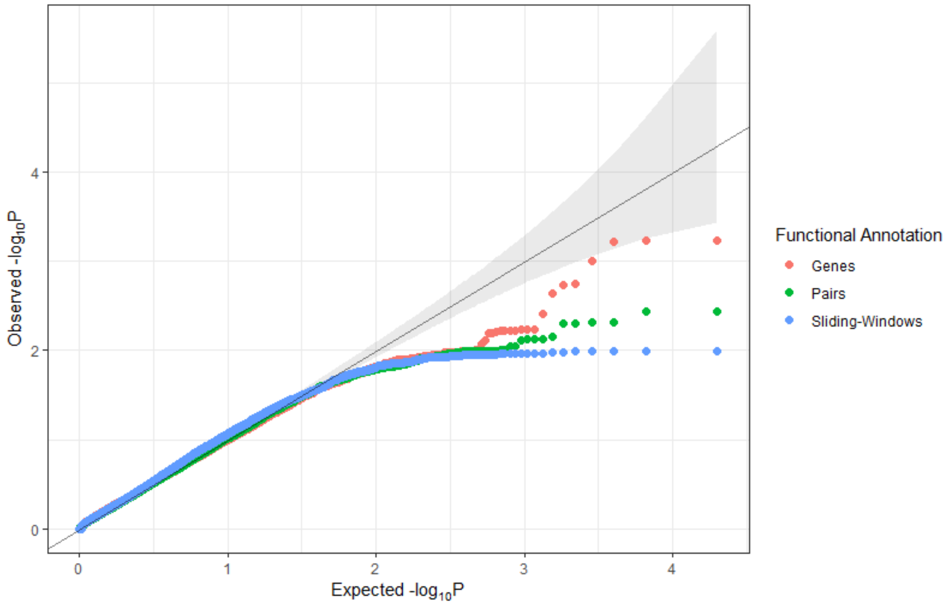

Figure S 9: Quantile-Quantile plots of ACAT-Combined bootstrap-based p-values for: RetroFun-RVS$_{Genes}$, RetroFun-RVS$_{Pairs}$, and RetroFun-RVS$_{Sliding-Windows}$. P-values were computed empirically based on 1,000 bootstraps for asymptotic ACAT-combined p-values between 0.01 and 0.001 and based on 10,000 bootstraps for asymptotic ACAT-combined p-values less than 0.001. Arbitrary values corresponding to 1/total number of bootstrap samples were given when empirical single-annotation p-values were 0.

#### 3.2 Power

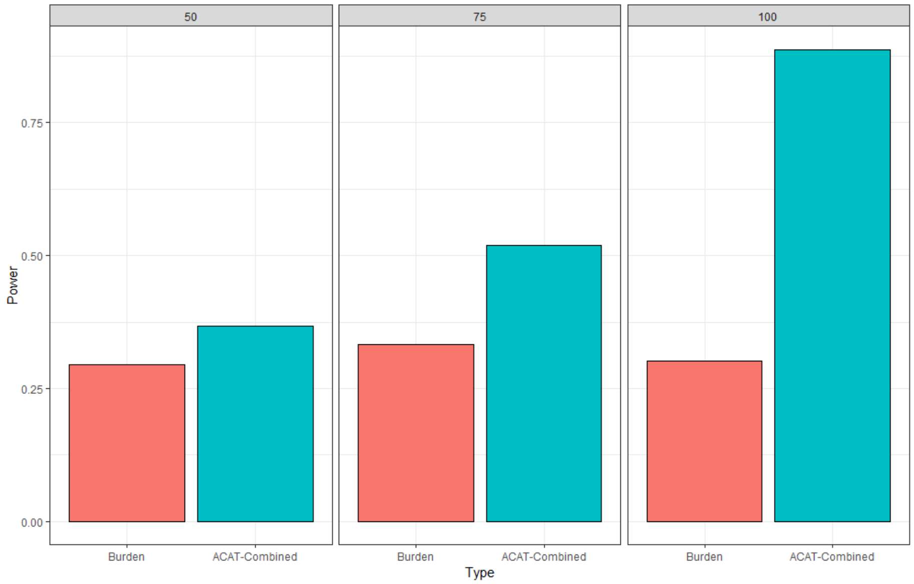

Figure S 10: Power evaluation of RetroFun-RVS under different scenarios for 2% risk variants considering only pedigrees of small to moderate size. Power at different proportions of risk variants within the CRH, between RetroFun-RVS$_{CRHs}$ with no functional annotation (Burden Original) and RetroFun-RVS$_{CRHs}$ including the four CRHs (ACAT-Combined). Power was evaluated on 1,000 replicates.

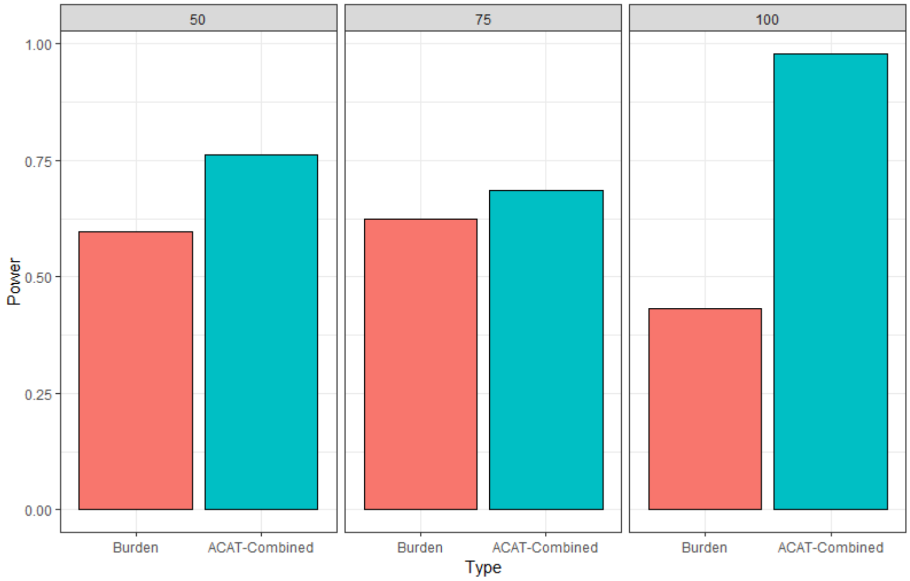

Figure S 11: Power evaluation of RetroFun-RVS under different scenarios for 1% risk variants. Power at different proportions of risk variants within the CRH, between RetroFun-RVS$_{CRHs}$ with no functional annotation (Burden Original) and RetroFun-RVS$_{CRHs}$ including the four CRHs (ACAT-Combined). As mentioned in the main manuscript, functional annotations with less than five families were removed from the analysis for ensuring a proper asymptotic behavior. Power was evaluated on 1,000 replicates. We noticed that the Burden Original statistic varies across 75% or 50% causal scenarios and 100% causal. Investigating this aspect, we found that this may due to simulation specificity, where in 100% causal scenario, statistic values are smaller compared to the two others scenarios (average statistic values of 2,748,881; 3,080,295 and 3,110,373, for 100%, 75% and 50% causal, respectively), while variances remain stable (average statistic values of 116,071; 113,284; 116,541). We hypothesize that could be due to preferential choice of small families in 100% causal compared to 75% causal or 50% causal (average Pearson correlation: 0.83). See Figure S17.

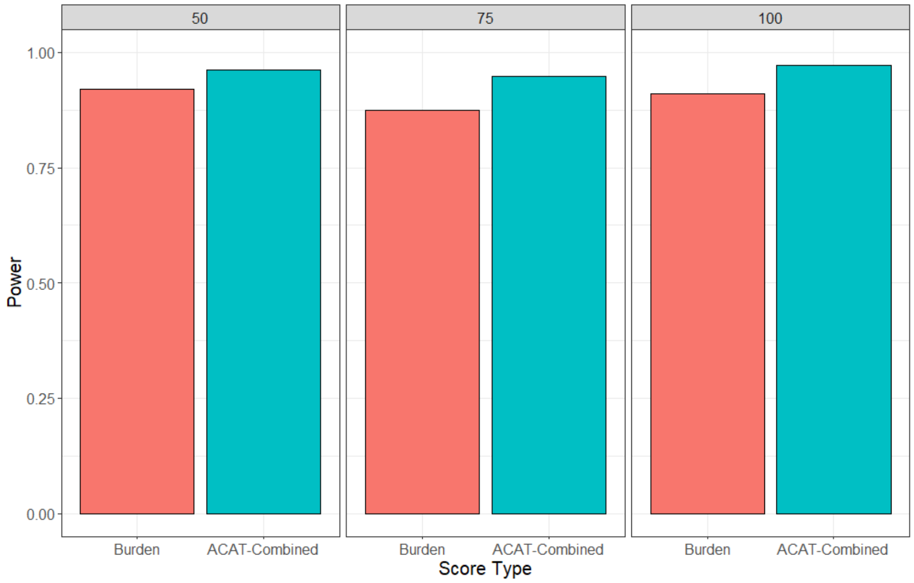

Figure S 12: Power evaluation of the original Burden test and ACAT-Combined p-values under different scenarios for 2% risk variants for RetroFun-RVS$_{Sliding-Windows}$. As mentioned in the main manuscript, functional annotations with less than five families were removed from the analysis for ensuring a proper asymptotic behavior. Power was evaluated on 1,000 replicates.

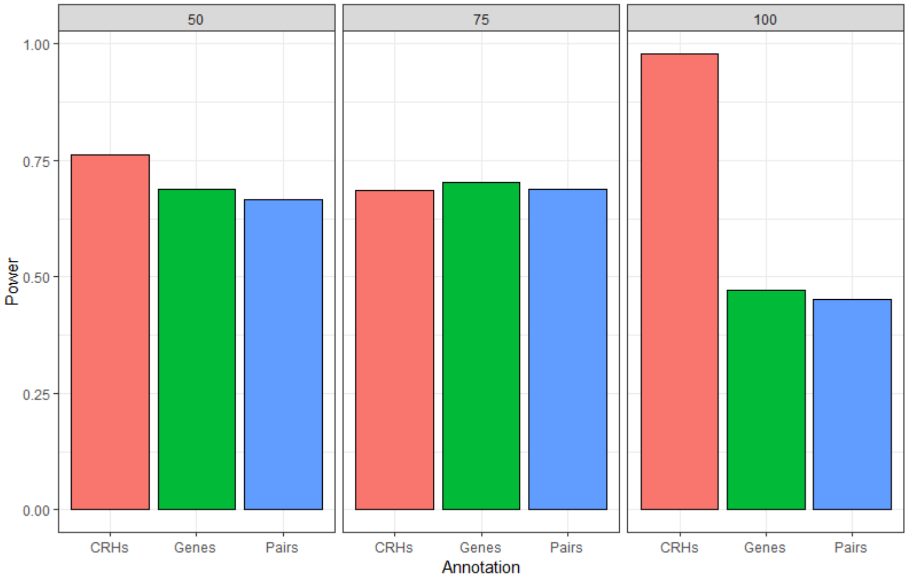

Figure S 13: Power evaluation of ACAT-Combined p-values under different scenarios for 1% risk variants for RetroFun-RVS$_{CRHs}$ (CRHs), RetroFun-RVS$_{Pairs}$ (G-E Pairs), RetroFun-RVS$_{Genes}$ (Genes), and RetroFun-RVS$_{Sliding-Window}$ (Sliding). As mentioned in the main manuscript, functional annotations with fewer than five families were removed from the analysis. Power was evaluated on 1,000 replicates.

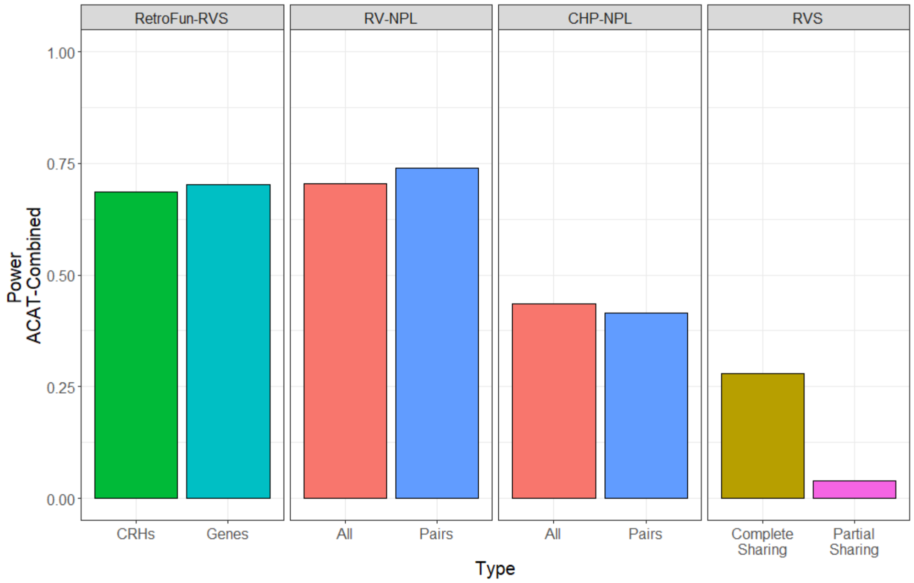

Figure S 14: Power at 75% risk variants within one CRH between RetroFun-RVS$_{CRHs}$ and other affected-only competiting methods for 1% causal variant. Here we included RetroFun-RV$_{genes}$ to micmic CHP-NPL procedure. As mentioned in the main manuscript, when considering RetroFun-RVS, functional annotations with less than five families were removed from the analysis for ensuring a proper asymptotic behavior. Power for RetroFun-RVS$_{CRHs}$ and RetroFun-RVS$_{Genes}$ was evaluated on 1,000 replicates while for RV-NPL and RVS we generated 200 replicates.

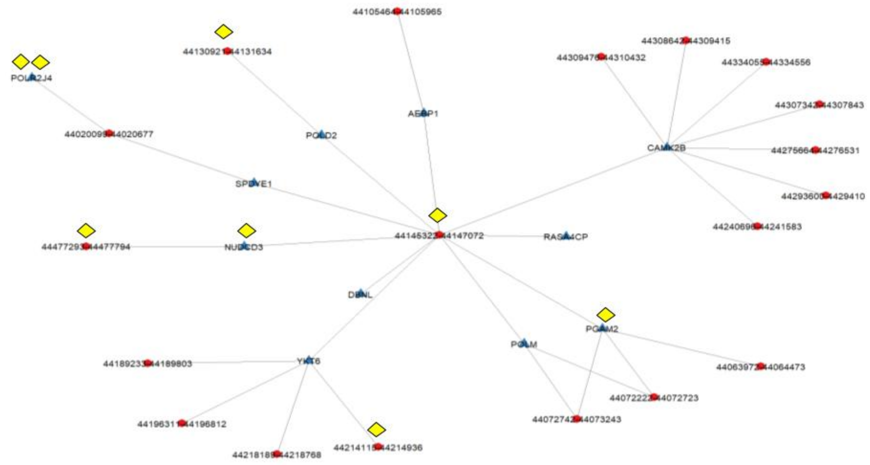

Figure S 15: Cis-Regulatory Hub on chromosome 7 highlighting 11 genes (blue triangles) and 19 enhancers (red circles). Rare variants are illustrated by yellow diamonds.

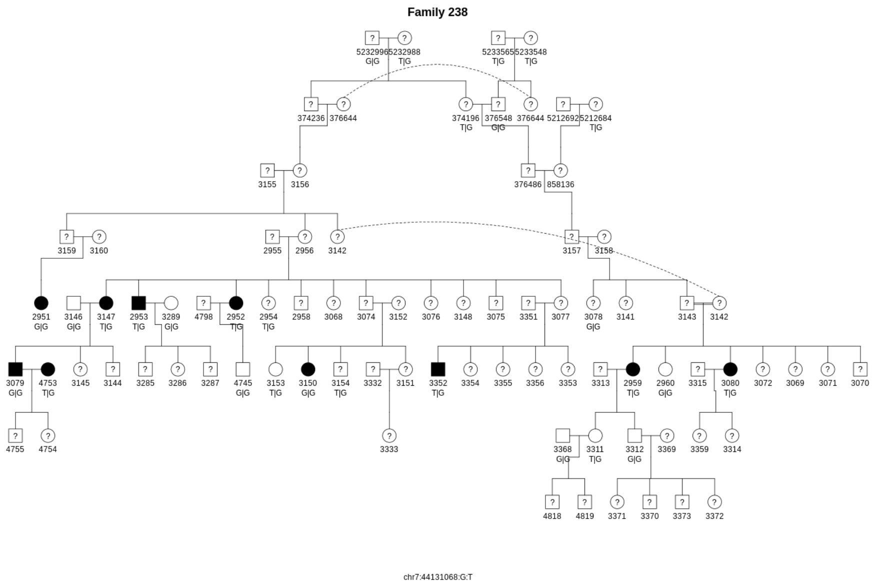

Figure S 16: Family with the rare T allele at SNV chr7:44131068 shared by the largest number of affected relatives within a cis-regulatory hub detected by RetroFun-RVS. Being a pedigree founder (no known parents), affected subject 4753 shares the T allele with other affected family members through unknown relations accounted for by the correction for cryptic relatedness based on the kinship among founders. Position is in GRCh38 coordinates.

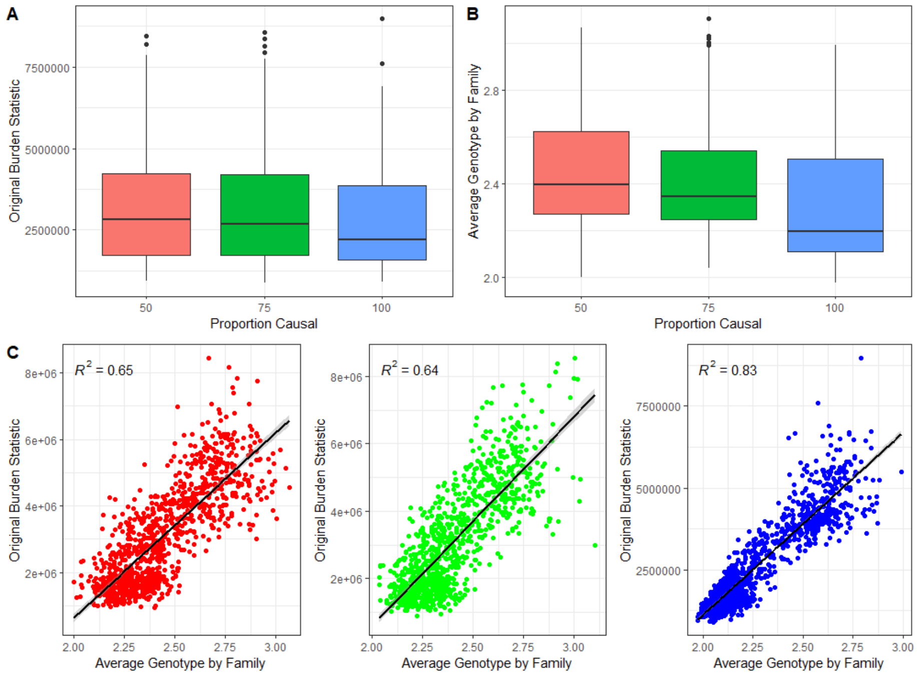

Figure S 17: Relationship between average genotype values by family and Burden Original statistic at 1% causal. (**A**) Boxplots across the 1,000 replicates of the Burden Original statistic for each scenario. (**B**) Boxplots across the 1,000 replicates of the average genotype value by family. (**C**) Regression line across the 1,000 replicates between the average genotype value by family and the Burden Original statistic.
